# EXOSC4 is recruited by histone H3 co-modified with K9me3 and acetylations to surveil non-coding transcription

**DOI:** 10.1101/2024.08.05.606680

**Authors:** Stephanie Stransky, Ronald Cutler, Jennifer Aguilan, Joseph D. DeAngelo, David Shechter, Simone Sidoli

## Abstract

Histones are hyper modified proteins that regulate chromatin accessibility and DNA readout. Co-existing post-translational modifications (PTMs) on histones affect interaction affinities of chromatin-associated proteins in ways that are still mostly unexplored. Here, we focus on the biological role of a specific histone code made of two PTMs with supposedly opposing biological functions, i.e. H3K9me3 marker of constitutive heterochromatin and H3K14ac benchmarking accessible chromatin. By applying multi-dimensional mass spectrometry, we demonstrated that EXOSC4 interacts with H3K9me3 + acetyls and affects post-transcriptional regulation. Specifically, EXOSC4 depletion leads to down-regulation of the RNA surveillance machinery and increased expression of non-coding transcripts, including anti-sense RNAs. Together, we elucidate the role of a co-modified histone tail and demonstrate its involvement in the RNA machinery and spurious transcription surveillance.

## Introduction

With the advancement of mass spectrometry and the analysis of protein post-translational modifications (PTMs), it has become clear that many mammalian proteins are hyper modified and they frequently carry co-existing PTMs. The histone protein family is maybe the clearest example, as they are modified with hundreds of different PTMs, many of them are still being discovered (*1*). About 25 years ago, Allis and colleagues proposed the existence of a “histone code” implying that co-existing modifications work as “keys” in a “lock” being protein complexes in charge of interpreting DNA readout (*2*). Since then, a few cross-talks between histone PTMs have been identified, the first example being the H3K9me3S10ph co-occurrence (*3*). H3K9me3 recruits the heterochromatin protein HP1, of which binding is impaired by the presence of H3S10ph, demonstrating that even well-characterized histone marks have a different impact on recruitment of the chromatin proteome when other modifications co-exist on the same histone sequence. Several publications showed that histone modifications with theoretically opposing biological functions frequently co-exist, e.g. silencing methylations with acetylations associated with open chromatin (*4–6*). Since methods like middle-down and top-down mass spectrometry have evolved to study histone tails and intact histones, respectively, many combinatorial histone PTM signatures have been found (*7, 8*). However, it is unclear which histone marks have a dominant vs recessive impact on transcriptional output, or whether the co-existence of antagonistic PTMs lead to a whole new biological function not associated with either of the individual ones.

Modifications such as H3K9me3K14ac are a puzzle for chromatin biologists, as H3K9me3 is arguably the best characterized marker of constitutive heterochromatin (*9–11*) while histone acetylation benchmarks accessible and transcribed chromatin (*12–14*). However, many studies have shown that this modification occupies 1-5% of most mammalian cells (*4–6*). Functionally, this histone code was shown to define chromatin regions (*15*), similarly to the bivalent domain in stem cells H3K4me3K27me3 (*16*). However, the following questions remain unanswered: (i) whether these chromatin domains regulate transcription of coding regions, (ii) which protein readers recognize these histone codes, (iii) whether this phenomenon is regulated in cells upon which signals. Therefore, identifying specific effector proteins recognizing these co-occupying marks is essential.

Studies have suggested that RNA surveillance has a central role in chromatin modification and gene expression (*17–19*). This quality control machinery starts at the chromatin level where DNA-associated RNAs are degraded to avoid genomic instability following nuclear and cytoplasmic regulation to control the titration of ncRNA and mRNA levels (*20*). One of the components of the surveillance system is the RNA exosome complex, which is a highly conserved multiprotein structure formed by exosome subunits (EXOSCs). This 3’-5’ exoribonuclease complex is localized in both the nucleus and the cytoplasm and plays a pivotal role in regulating gene expression as well as in maintaining RNA homeostasis, ranging from the degradation of aberrant or unwanted transcripts to the processing of precursor RNAs (*21–23*). The core of the RNA exosome is a barrel-shaped catalytically inactive structure formed by nine subunits. It comprises the ‘cap’ structure (EXOSC1-EXOSC3), which contribute to the RNA binding and the ‘ring’ complex (EXOSC4-EXOSC9), which is essential for the RNA processing and degradation (*23, 24*). The interplay between these subunits allows the exosome to recognize and process a wide range of RNA substrates, including messenger RNA (mRNA), ribosomal RNA (rRNA), transfer RNA (tRNA), and small nuclear RNA (snRNA) (*21*). Loss-of-function of the exosome complex triggered by mutations or depletion of its subunits can cause alterations in many biological mechanisms (*22*). EXOSC4, also known as RRP41, is a non-catalytic component of the RNA exosome complex. The dynamic nature of the function of EXOSC4 underscores its relevance in diverse cellular processes. Studies have implicated dysregulated EXOSC4 expression in reducing pancreatic and liver cancer cell growth and triggering cell death through apoptosis (*25, 26*). In addition, it was previously shown that EXOSC4 overexpression increases the proliferation (*27*). Although it was demonstrated that EXOSC4 has a potential role in tumorigenesis, its molecular mechanisms are still poorly elucidated.

In this paper, we show that our biological model where we induced anomalous decondensed heterochromatin, i.e. via HDAC inhibition with sodium butyrate treatment, are decorated by a histone tail co-modified with H3K9me3 and acetylations which specifically recruits the nuclear exosome via interaction with EXOSC4. We further demonstrate that EXOSC4 and its protein interactors surveil spurious transcription, contributing to maintaining low levels of non-coding transcripts and, therefore minimizing genomic instability. Understanding the role of combinatorial histone PTMs and their specific readers during anomalous chromatin decondensation will provide insights into the intricate network of proteins that can be targeted for several disease therapies.

## Results

### Decondensed heterochromatin is enriched by co-modified histones

Our initial goal was to understand co-occupancy of histone modifications on hyperacetylated and decondensed chromatin. To modulate chromatin decondensation, we treated cells with NaBut, a histone deacetylase inhibitor (HDACi) that causes hyperacetylation of histone H3 and H4 leading to chromatin remodeling towards an open and transcriptionally state (*28*). Overall, the global levels of mono, di, or trimethylated histone H3 peptides did not differ between NaBut-treated and non-treated cells, but there was an increase in the levels of acetylation (**Fig. 1A**).

**Fig. 1.**
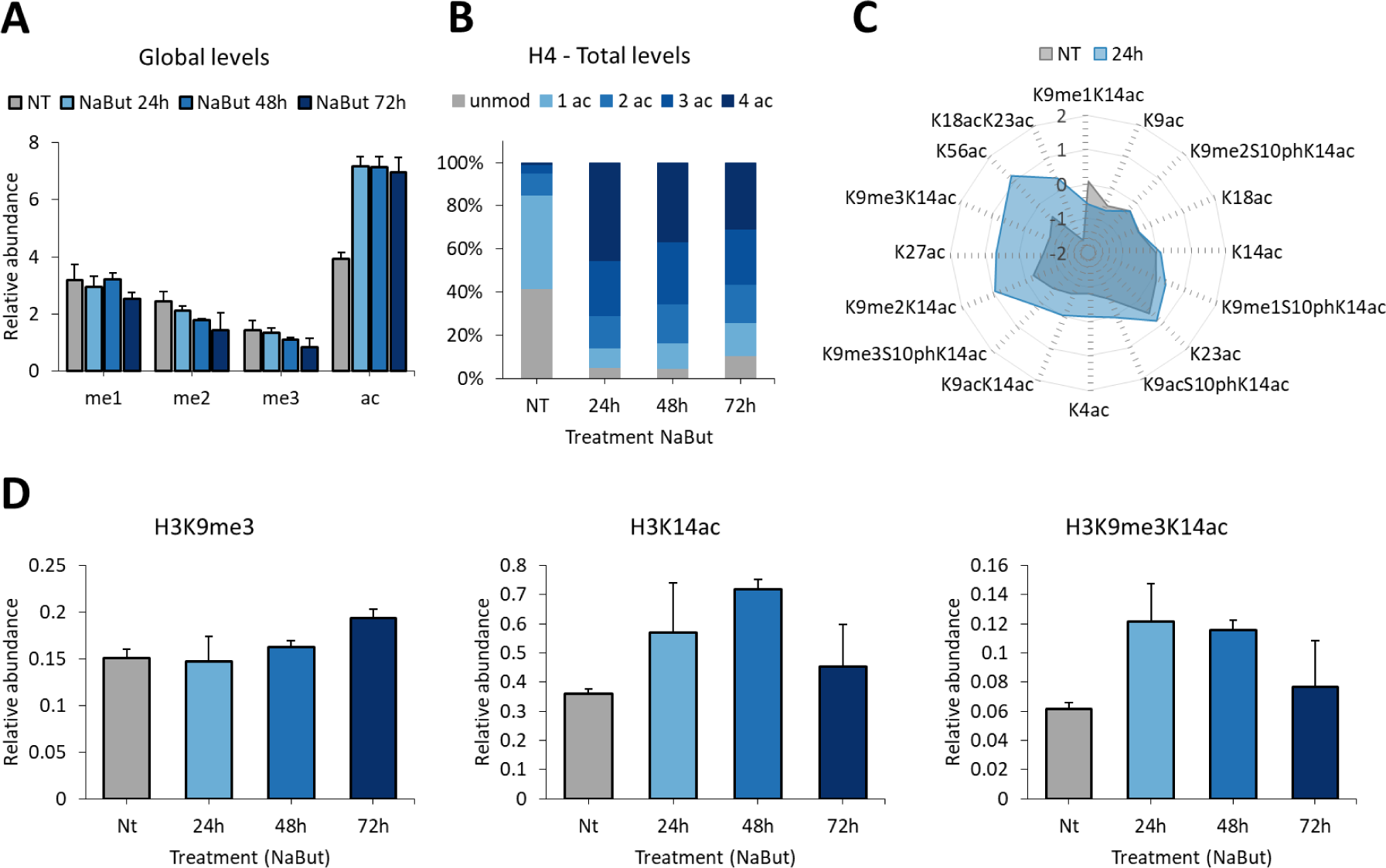
Histones co-modified with K9me3 and K14ac benchmark regions of decondensed heterochromatin. HepG2/C3A flat cells were treated with 20 mM of NaBut for 24, 48 and 72 h. Histones were extracted and analyzed by bottom-up mass spectrometry. **(A)** Global levels of histone peptides containing 1, 2 or 3 methylations (me1, me2, me3 respectively) or acetylations (ac). **(B)** Total levels of histone H4 peptides containing multiple acetylations (1ac, 2ac, 3ac, 4ac). Unmod, unmodified peptide. **(C)** Combinatorial histone marks enriched in NaBut- (24h, blue) and non-treated cells (NT, grey). **(D)** The bar graphs show the relative abundance of histone H3K9me3, H3K14ac and H3K9me3K14ac. Data are represented as means ± SEM.

More specifically, there was an accumulation of hyperacetylated histones, containing 2, 3 and 4 acetyl groups, following treatment for 24, 48 or 72 h (**Fig. 1B**).

The analysis of histone PTMs by bottom-up mass spectrometry allows the identification and quantification of single and co-existing modifications (*29*) (**Supplementary Table 1**). We detected on NaBut-treated cells histones singly modified with acetylation but also co-modified with methylations (me1/me2/me3), acetylations (ac) and phosphorylations (ph) (**Fig. 1C**).

Among the co-modified histone H3 peptides, K9me2/3K14ac and K9me3S10phK14ac were the main upregulated marks in treated cells. Although high levels of K9me3K14ac were detected, the single mark K9me3 was not regulated (**Fig. 1D**), suggesting that NaBut-induced chromatin decondensation is not correlated with loss of silencing histone PTMs but with decoration of silenced chromatin domains with PTMs that decorate open chromatin.

### Nuclear EXOSC4 was enriched by histone H3 peptides co-modified with the silencing mark H3K9me3 and acetylations

To investigate the proteins binding to histones co-modified with H3K9me3 and acetylations, we performed a peptide pull-down experiment combined with mass spectrometry (**Fig. 2A**). In this assay, three synthetic histone H3 peptides carrying the PTMs of interested and conjugated to biotin were chemically synthesized: (i) K9me3, (ii) K14acK18acK23ac – 3 acetyls, and (iii) K9me3K14acK18acK23ac – hybrid. The peptides were immobilized onto streptavidin beads, incubated with soluble nuclear proteins (SNF) or with proteins that bind to chromatin (CHR) and then the interactome of each peptide was analyzed by mass spectrometry (**Supplementary Table 2**). As a quality control of the assay, we identified known chromatin readers, such as chromodomains that bind to histone methylation and bromodomains that bind to hyperacetylated histones (**Fig. S1**). Our screening of the hybrid peptide interactome revealed proteins involved in the regulation of RNA processes and chromatin structure (**Fig. 2B**). We then compared the interactome of the hybrid peptide with the K9me3 and 3 acetyls peptides and EXOSC4 was detected as one of the most enriched proteins in the pull-down with the hybrid modified peptide (**Fig. 2C**). We showed that this binding is highly specific, as no interaction with the other histone H3 peptides was observed (**Fig. 2D**). We also demonstrated that hyperacetylation induced by NaBut leads to increased levels of EXOSC4 (**Fig. 2E**).

**Fig. 2.**
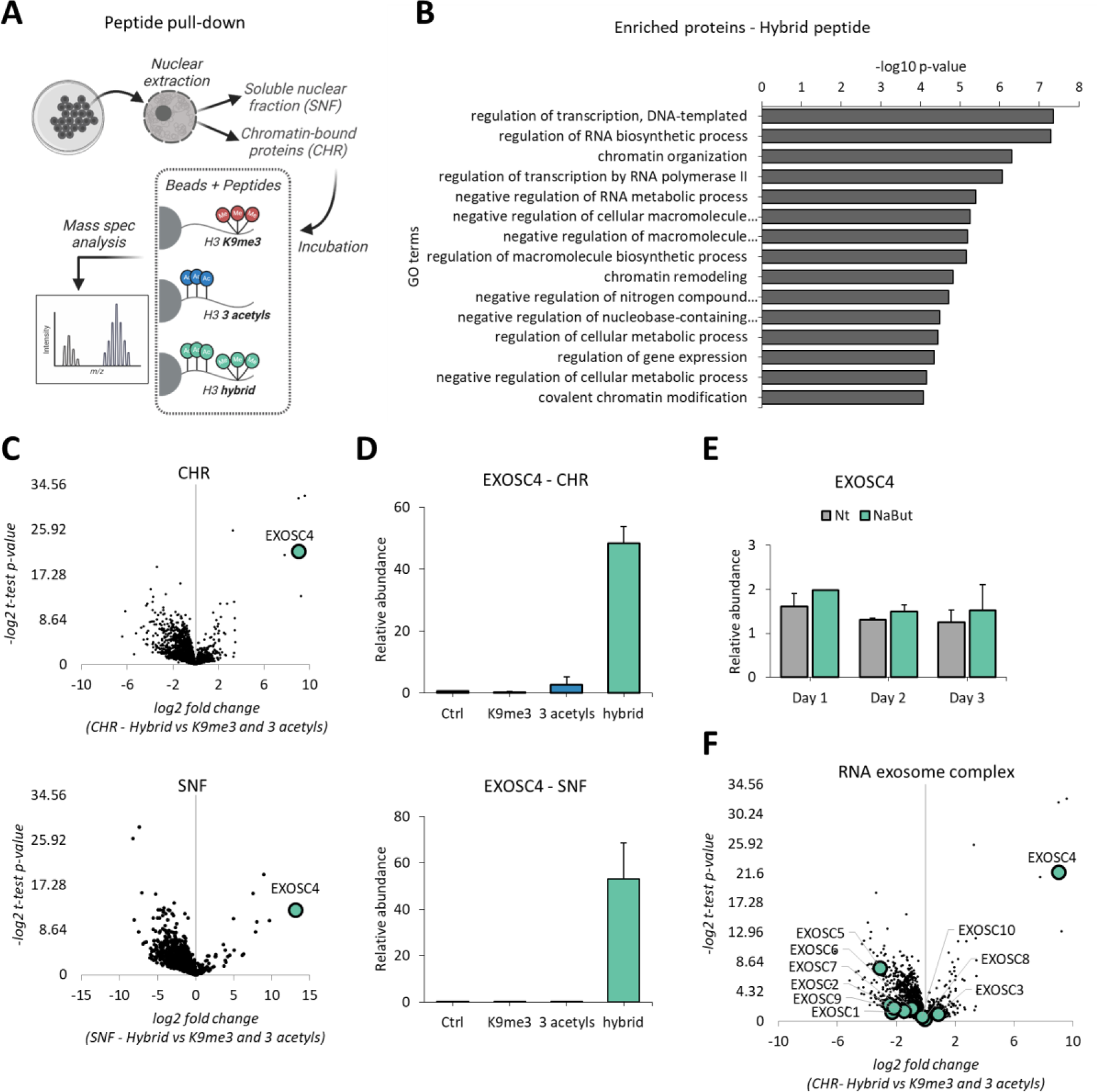
EXOSC4 is a reader of the combinatorial mark K9me3 + 3 acetyls. **(A)** Scheme of peptide pull-down experimental approach. Created using BioRender.com. **(B)** Gene ontology (GO) enrichment analysis of up-regulated proteins bound to the synthetic hybrid peptide. Functional annotation was obtained using GOrilla (*30*). **(C)** The volcano plot shows the fold change relative abundance of chromatin (CHR) and soluble nuclear (SNF) proteins bound to the hybrid histone mark versus proteins bound to the K9me3 and 3 acetyls marks. Highlighted bubble indicates EXOSC4. **(D)** The bar graph shows the EXOSC4 pulled down by the K9me3, 3 acetyls, and hybrid modified histone peptides in the chromatin (CHR) and soluble nuclear fractions (SNF). **(E)** Bar graph showing the EXOSC4 relative abundance in NaBut-treated cells compared to non-treated cells (Nt). Data are represented as means ± SEM. **(F)** The volcano plot shows the fold change relative abundance of chromatin (CHR) proteins bound to the hybrid histone mark versus proteins bound to the K9me3 and 3 acetyls marks. Highlighted bubble indicates the components of the RNA exosome complex.

Exosome component 4 (EXOSC4, RRP41) is one of the RNA exosome complex subunits and participates in cellular RNA processing and degradation events (*27*). Notably, even though we have detected all the components from the RNA exosome complex in our proteomics dataset, only EXOSC4 was significantly enriched in the hybrid peptide (**Fig. 2F**).

### Interactors of EXOSC4 are involved in RNA processing pathways

To better understand the role of EXOSC4 in domains of decondensed heterochromatin and its potential interactors, we performed a pull-down using the protein as bait. For this assay, nuclear and cytoplasmic extracts were incubated with immobilized EXOSC4, and the co-precipitated proteins were detected by mass spectrometry (**Fig. 3A**) (**Supplementary Table 3**). We confirmed that proteins enriched by EXOSC4 are also part of different RNA processing pathways, both in the nucleus (**Fig. 3B**) and cytoplasm (**Fig. 3D**). In fact, proteins that regulate mRNA metabolism, RNA transport and the structure of nucleosomes are found to be equally enriched (**Fig. 3C and E**). On the other hand, terms such as protein-DNA complex assembly and chromatin remodeling are exclusively found in the nucleus fraction (**Fig. 3C**) and ncRNA (non-coding RNA) processing and regulation of mRNA stability are found only in the cytoplasm (**Fig. 3E**). These findings show the importance of EXOSC4 and its interactors in regulating RNA surveillance pathways and modulating nucleosome and chromatin structure.

**Fig. 3.**
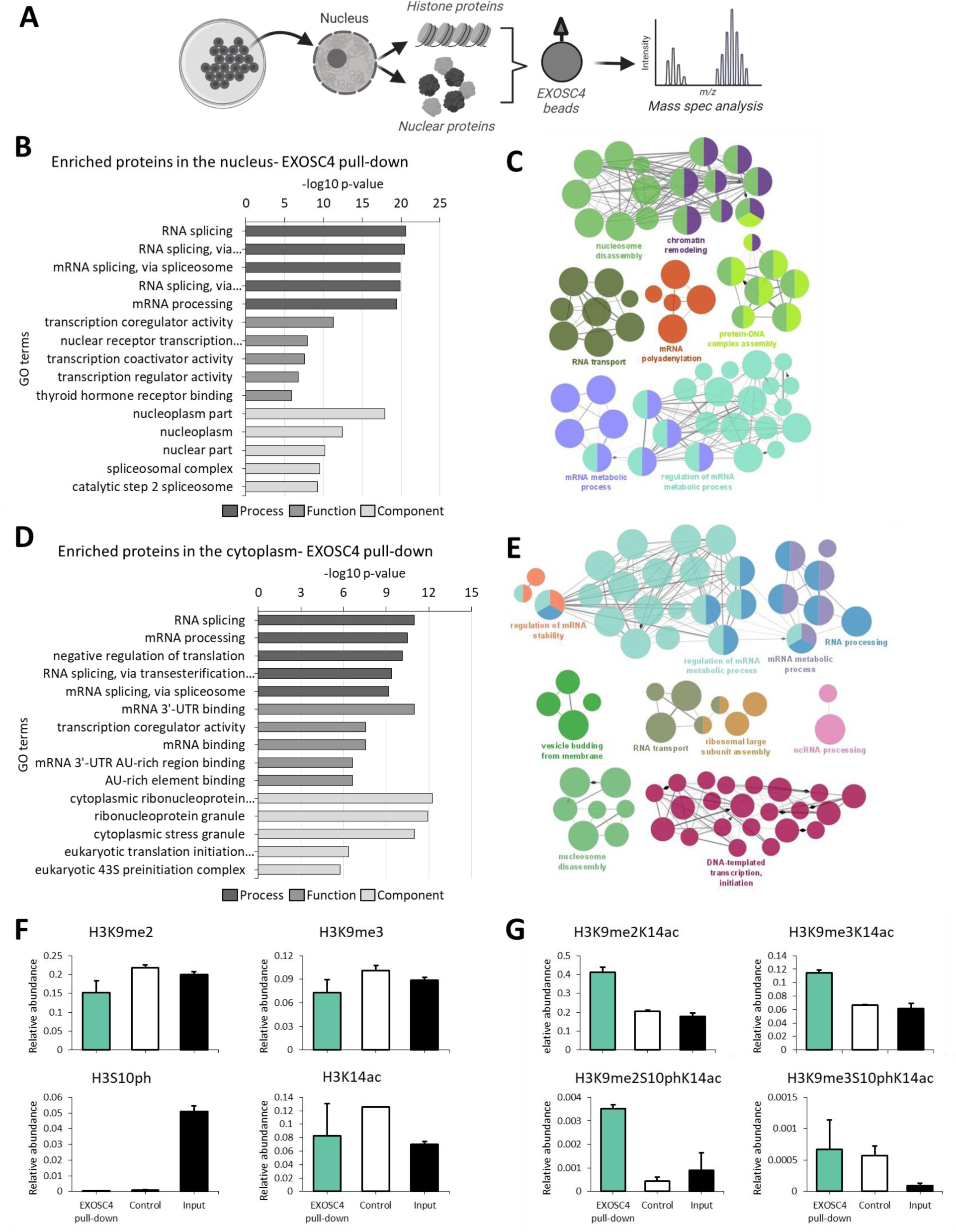
Interactors of EXOSC4 regulate RNA processing pathways. **(A)** Scheme of EXOSC4 pull-down experimental approach Created using BioRender.com. **(B)** GO enrichment analysis of the nuclear proteins enriched by EXOSC4 beads. **(C)** Clustered network of 200 enriched nuclear proteins by EXOSC4 beads. Clusters reflect connected portions of the network and correspond to functional categories of the proteins, with terms as nodes. The network was constructed by ClueGo (*31*), a Cytoscape plug-in. **(D)** GO enrichment analysis of the cytoplasmic proteins enriched by EXOSC4 beads. **(E)** Clustered network of 200 cytoplasmic proteins enriched by EXOSC4 beads. **(F)** Relative abundance of histone H3 peptides containing single marks that are recruited by EXOSC4. **(G)** Relative abundance of histone H3 peptides containing combinatorial marks recruited by EXOSC4. Data are represented as means ± SEM. Control, streptavidin beads.

As EXOSC4 was found to be specifically enriched by the combinatorial mark H3K9me3K14acK18acK23ac, we next investigated the interaction between other histone marks and EXOSC4 by using the pull-down approach, as a reverse validation. EXOSC4 beads, as well as streptavidin beads (Control), were incubated with histones extracted from HepG2/C3A cells, and the enriched histone peptides were analyzed by mass spectrometry (**Supplementary Table 4**). The input material was analyzed and used as a reference. Repressive and activating marks, such as H3K9me3 and H3K14ac respectively, as well as phosphorylation marks were not specifically enriched in the EXOSC4 pull-down compared to controls (**Fig. 3F**). However, the enrichment of combinatorial marks, such as H3K9me2K14ac, H3K9me3K14ac, and H3K9me2S10phK14ac, appears to be specific (**Fig. 3G**) when compared to control and input, confirming the preference of EXOSC4 for histone tails co-modified with methylation and acetylation.

### EXOSC4 is a key component of the RNA surveillance machinery

To confirm the role of EXOSC4 in RNA processing pathways, we generated HepG2/C3A cells stably expressing dCas9-KRAB-MeCP2 (**Fig 4A**) (*32*). We evaluated the cell fitness following gRNA transduction and, although we observed more cells in the supernatant compared to cells transduced with a scrambled gRNA (Scr), knocked-down cells (KD) remained viable (**Fig. 4B**). Western blot (**Fig 4C**), mass spectrometry (**Fig 4D**), and RNA sequencing (RNA-seq, **Fig 4E**) analyses confirmed that efficient EXOSC4 depletion was achieved (**Fig. S2**). In fact, depletion of EXOSC4 led to co-depletion of the other exosome components (**Fig. 4F and G**), as previously observed by (*33*). The gene ontology enrichment analysis revealed that EXOSC4 depletion leads to downregulation of the RNA surveillance machinery and non-coding pathways (**Fig. 4H**). RNA surveillance machinery refers to cellular processes that detect and degrade aberrant and potentially harmful RNAs, ensuring the quality and integrity of the transcriptome (*22*). Although upregulated proteins revealed a poor enrichment, they showed to be related to cell death and vesicle-mediated transport (**Fig. 4H**).

**Fig. 4.**
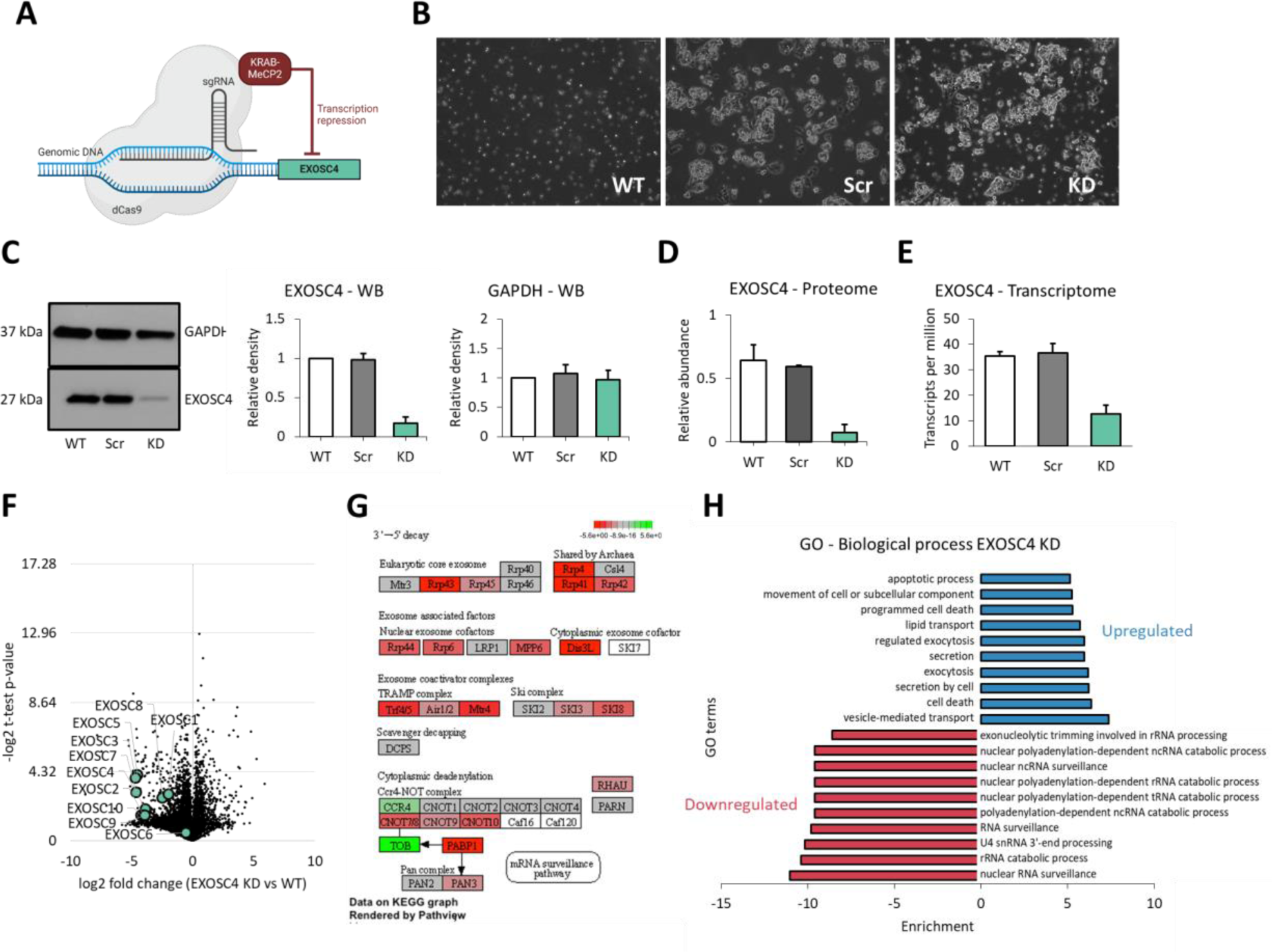
EXOSC4 is involved in RNA surveillance processes. **(A)** Scheme of the CRISPRi protocol. The dcas9-KRAB-MeCP2 expression vector was used to repress EXOSC4. **(B)** HepG2/C3A cell morphology following non-related (Scr, scrambled) or EXOSC4 (KD, knockdown) gRNA stable transduction. Wild-type (WT) represents non-transduced cells. **(C)** Western Blot analysis of HepG2/C3A cell lysates after EXOSC4 knockdown. The western blot membrane was probed with anti-EXOSC4 or anti-GAPDH antibody (loading control). The bands were quantified and the data are represented as means ± SEM. **(D)** The bar graph shows the levels of EXOSC4 following transduction with Scr or EXOSC4. The proteome of the cells was analyzed by mass spectrometry and the transcriptome by RNA-seq **(E)**. Data are represented as means ± SEM. **(F)** The volcano plot shows regulation of other RNA exosome complex (EXOSCs) components on knockdown cells (KD) at the protein level. **(G)** Regulation of other RNA exosome complex (EXOSCs) components on knockdown cells (KD) at the RNA level. KEGG pathway map was acquired from Pathview software (*34*). **(H)** Gene ontology (GO) analysis of biological processes that are up and downregulated in KD cells. Functional annotation was obtained using GOrilla (*30*).

H3K9me3 is known to decorate regions of constitutive heterochromatin by repressing repetitive elements, a.k.a. non-coding sequences that pose a serious challenge to genome stability (*35, 36*). Even though non-coding RNAs (ncRNAs) lack protein-coding potential, they play pivotal roles in diverse cellular processes, including chromatin remodeling and transcriptional regulation(*37*). Notably, we showed here that NaBut treatment induced decondensation of heterochromatic domains and accumulation of the combinatorial mark H3K9me3K14ac (**Fig. 1D**). To demonstrate that EXOSC4 KD regulates the transcriptome with its specificity, i.e. it is not the equivalent than an overall chromatin decondensation stress, we compared the transcriptomics results of NaBut treated cells with the results of the EXOSC4 KD experiment. Data was split between protein-coding transcripts, non-coding and pseudogenes (**Fig. 5A-B upper panel**). The impact of EXOSC4 KD on the levels of transcripts was not nearly as dramatic as by using NaBut (y-axis of **Fig. 5A-C upper and lower panels**). However, only EXOSC4 KD cells showed a significantly asymmetric shift in terms of non-coding RNA levels (**Fig. 5B lower panel**). Depletion of EXOSC4 led to a higher abundance of more than 5,000 long non-coding RNAs (lncRNAs). To further prove that the RNA levels regulated during NaBut treatment and EXOSC4 KD are different, we performed a correlation analysis (**Fig. 5D**), which was poor substantiating our hypothesis that EXOSC4 KD has a significantly different effect on RNA levels compared to overall chromatin decondensation stress. We observed some similarity in regulation among the protein-coding transcripts, but nearly no correlation was observed for the non-coding and pseudogene transcripts.

**Fig. 5.**
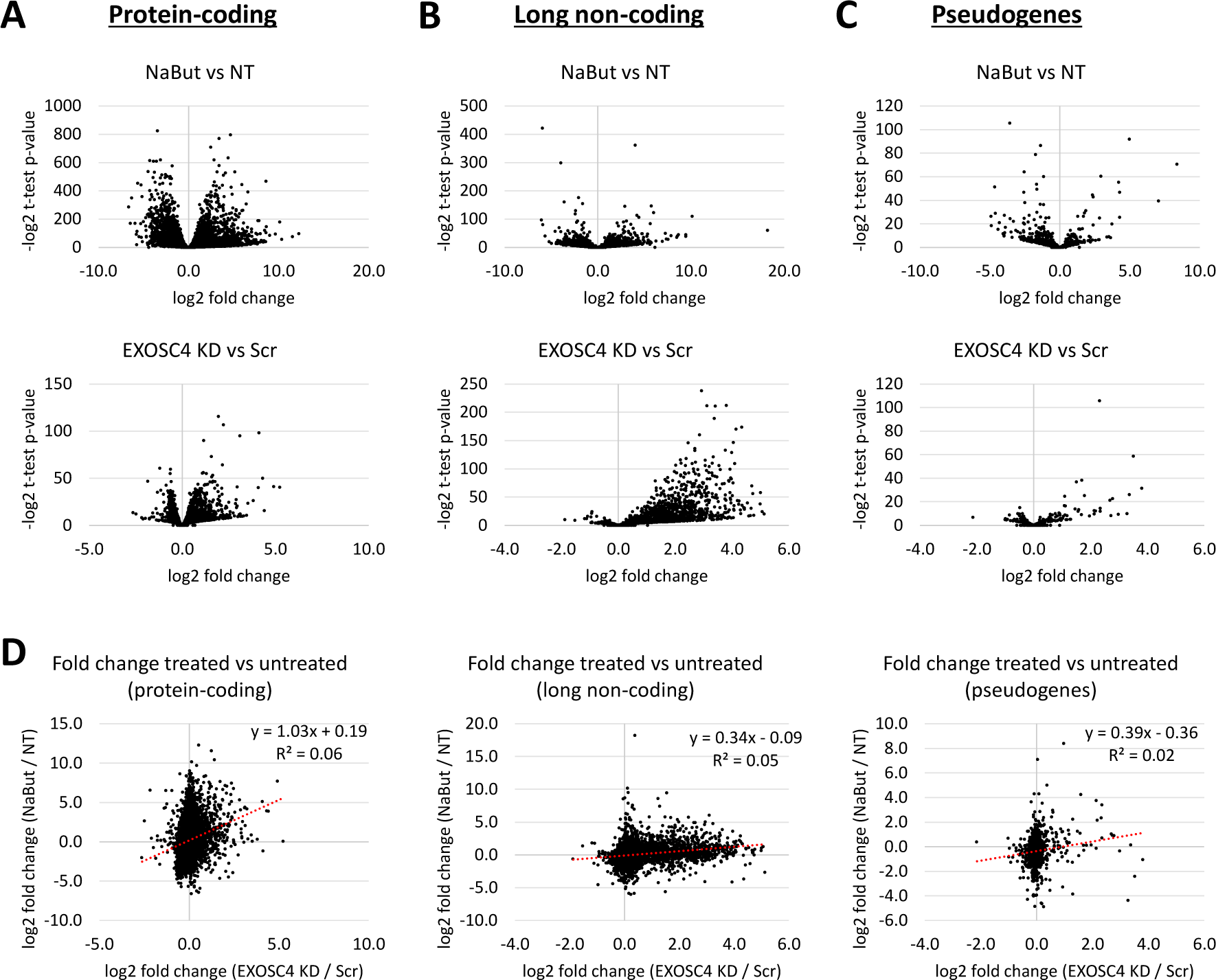
EXOSC4 regulates the expression of non-coding transcripts. Volcano plots showing the RNA levels of **(A)** protein coding and **(B)** non-coding transcripts and **(C)** pseudogenes in NaBut-treated and EXOSC4 KD cells. **(D)** Correlation of protein coding, non-coding transcripts and pseudogenes between NaBut-treated and EXOSC4 KD cells. KD knockdown; Scr, scrambled gRNA; NaBut, 20mM sodium butyrate; NT, non-treated.

Finally, we put in perspective the number of transcripts regulated by the absence of EXOSC4. Specifically, among the transcripts analyzed in KD cells, 25% were lncRNAs (**Fig. 6A**). Even though protein-coding transcripts were the most abundant biotype detected, lncRNAs were the transcripts with the highest average fold change, followed by small nucleolar RNAs (snoRNAs) (**Fig. 6B**). Among lncRNA biotypes, we identified antisense transcripts. These RNAs that are complementary to other endogenous RNAs, have been shown to regulate the neighbor genes and play important roles in the regulation of gene expression (*38, 39*). Interestingly, our data show a strong enrichment of antisense transcripts in EXOSC4 KD cells (**Fig. 6B-D**), while protein-coding genes were more or less equivalent in terms of overall fold change (**Fig. 6B**). Importantly, the top regulated antisense transcripts in EXOSC4 KD cells present an inverse correlation with their corresponding sense coding genes (**Fig. 6E**), suggesting that they are involved in post-transcriptionally regulating their respective coding transcripts.

**Fig. 6.**
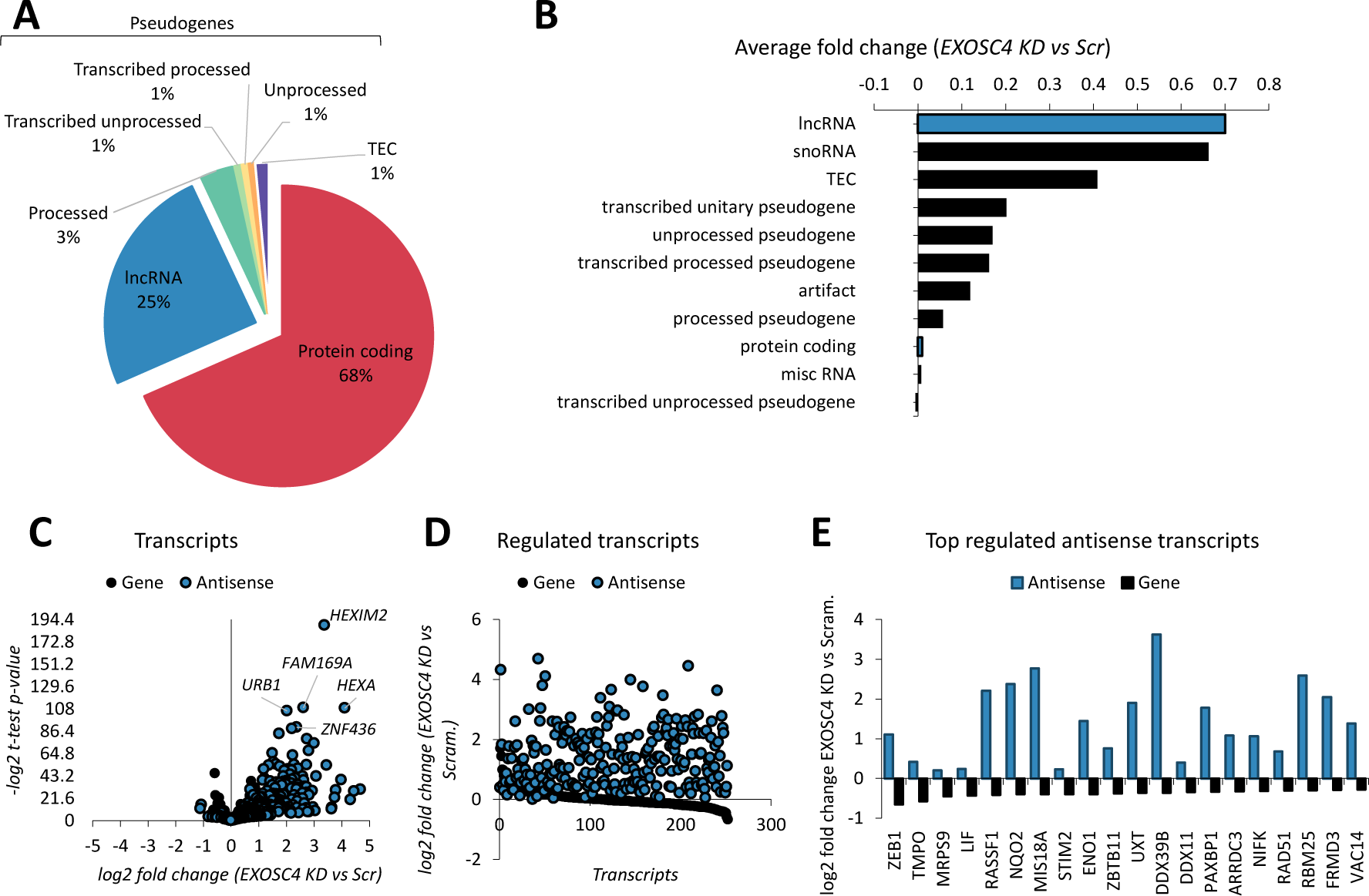
EXOSC4 KD cells are enriched in non-coding transcripts and anti-sense RNA. **(A)** Distribution of biotype representation for EXOSC4 KD cells. **(B)** Bar graph showing the average fold change per transcript biotype from EXOSC4 KD cells compared to Scr. Highlighted bars represent lncRNAs and protein-coding transcripts. **(C)** Volcano plot showing regulation of antisense RNAs in EXOSC4 KD cells compared to Scr. **(D)** Enrichment of protein-coding (gene) and anti-sense RNAs in EXOSC4 KD compared to Scr. **(E)** Expression levels of the top regulated antisense and their corresponding sense RNAs (gene) in EXOSC4 KD cells.

## Discussion

Over the past decades, research has revealed that histone PTMs do not act in isolation but rather form combinatorial patterns, resulting in a “histone code” that dictates gene expression and chromatin-related processes (*7, 40, 41*). Crosstalk between different histone modifications further amplifies the complexity of combinatorial histone marks, as certain modifications can either enhance or antagonize the function of others, leading to context-dependent outcomes in DNA readout (*42*). Although the biological relevance of co-existing PTMs is acknowledged, the mechanisms by which the combinatorial nature of these marks are incorporated into a biological pathway are not well characterized. Here, we investigated functional implications of combinatorial histone modifications under cellular stress and their role in chromatin instability.

Combinatorial histone modifications, such as H3K9me3 combined with acetylation, remain largely unexplored not only in their presence but also in their specific biological functions. We propose that heterochromatic domains that are decondensed upon cell stress, are decorated by this co-existing PTM, which has EXOSC4 as one of its readers. These complex modifications may represent an evolved cellular strategy to manage the transcriptional noise that becomes increasingly prevalent with different human conditions.

The establishment and maintenance of combinatorial histone codes are governed by a diverse array of histone modifying enzymes, chromatin remodelers, and reader proteins (*2*). The recruitment of specific proteins like EXOSC4 to regions marked by H3K9me3+acetyls underscores an adaptive response where cells have likely evolved mechanisms to specifically recognize and respond to these combinatorial modifications. We have shown that EXOSC4 and its interactors are part of the RNA surveillance machinery and play a key role in maintaining low levels of non-coding transcription. This specificity suggests a refined level of gene regulation aimed at controlling or mitigating the detrimental effects of spurious, cryptic transcription that has been shown to increase with age. For instance, studies have demonstrated that aging is associated with a loss of transcriptional fidelity and an increase in the transcription of normally repressed genomic regions, contributing to genomic instability and cellular senescence (*43, 44*).

Thought as transcriptional noise for a long time, non-coding RNAs (ncRNAs) are recognized now for mediating highly specific processes, such as regulation of gene expression, guiding RNA modifications or promoting posttranscriptional events (*36*). Recent findings revealed a widespread involvement of lncRNAs in the epigenetic machinery to control chromatin organization and gene expression. In fact, these RNAs have the capability to directly engage with enzymes responsible for modifying histones and DNA (*45*). Our findings that disruption in normal EXOSC4 function leads to an increase in antisense RNA production highlights the critical role of RNA surveillance mechanisms in maintaining genomic stability, especially under stress conditions that induce heterochromatin decondensation. This regulatory interplay becomes increasingly relevant specifically in aging cells, where the maintenance of genomic integrity is challenged, and the control over spurious transcription becomes crucial for cellular homeostasis.

The complex interplay between non-coding RNAs (ncRNAs) and chromatin modifications presents a significant frontier in our understanding of gene regulation. As we continue to uncover the extensive landscape of ncRNAs generated under stress conditions, the biological roles of these RNAs remain an intriguing puzzle. The upregulation of antisense and other non-coding RNAs in response to chromatin decondensation suggests that these molecules could play critical regulatory roles, potentially influencing gene expression, chromatin structure, and cellular phenotype. However, much work remains to elucidate these functions. Systematic studies are required to map the interactions between ncRNAs and the chromatin machinery, determine their impact on gene regulation, and understand their contribution to cellular homeostasis and disease.

Additionally, the potential therapeutic implications of targeting proteins that specifically recognize and bind to combinatorial histone modifications represent a promising area of research. For instance, disrupting the interaction between chromatin readers like EXOSC4 and their specific histone marks could be a strategy to selectively target cancer cells or clear senescent cells in aging tissues. The ability to modulate these interactions could lead to therapies that minimize the harmful effects of uncontrolled cell growth or the systemic effects of senescent cells (*46–48*). The development of therapies that can specifically inhibit these interactions may provide a targeted approach to treating diseases associated with chromatin misregulation, offering hope for interventions that could delay aging processes or improve cancer treatment outcomes.

In conclusion, our investigation not only extends the known repertoire of histone modifications but also suggests that the combinatorial nature of these modifications may encode a complex set of biological instructions. The recognition of unique histone modifications by specific proteins like EXOSC4 may be part of a broader cellular strategy to cope with the increased transcriptional noise observed in aging and different diseases. By further elucidating these relationships, future research can potentially identify novel targets for mitigating chromatin dysregulation.

## Materials and Methods

### Cell culture and treatment

HepG2/C3A flat cells (ATCC, CRL-10741) were maintained in Dulbecco’s Modified Eagle’s Medium (DMEM) supplemented with 10% Fetal Bovine Serum (FBS), 1% Non-Essential Amino Acids, 1% GlutaMAX, and 0.5% Penicillin/Streptomycin (all Corning) in a controlled atmosphere (5% CO_2_ incubator at 37°C). Flat cells were cultured until reach 80% confluency and then trypsinized using 0.5% Trypsin/EDTA diluted 1:1 in Hanks’ Balanced Salt Solution (HBSS, Corning) for 5 min, followed by centrifugation at 270 g for 5 min. Cells were counted using the Corning Cell Counter and CytoSMART Cloud App (Corning).

For pull-down experiments, cells (1 x 10^6^ cells/well) were plated in a 10 cm dish and cultured until reached 80% confluency. Cells were then collected using a cell scraper, counted and 2 x 10^6^ cells were split into tubes for replicates. Cell pellets were washed with HBSS and centrifuged for 5 min at 270 g. The supernatant was discarded and the dry pellets were stored in the -80°C freezer until further processing.

For the epigenetics drug treatment, cells (0.5 x 10^6^ cells/well) were plated in a 6-well plate and treated with 20 mM sodium butyrate (NaBut) for 24, 48 and 72 h. After each time point, the supernatant was discarded, cells were washed with HBSS, collected using a cell scraper and centrifuged for 5 min at 270 g. The supernatant was discarded and the dry pellet was stored in the -80°C freezer until further processing.

### CRISPRi

The dCas9-KRAB-MeCP2 expression vector (#122205) (*49*) and the EXOSC4-targeting gRNA vector (pXPR, #96925) (*50*) were purchased from Addgene. Custom gRNA sequences (**Supplementary Table 5**) derived from CRISPick were cloned as described previously (*51*). Lentiviral particles containing expression vectors were produced in HEK293T cells using calcium phosphate transfection. Transduction of HepG2/C3A cells was accomplished by combining lentivirus with media containing 4 µg/mL polybrene (Santa Cruz) and centrifuging at 30°C for 90 minutes at 500 g followed by incubation for 24 h in a controlled atmosphere (5% CO_2_ incubator at 37°C). Following the 24-h incubation, lentiviral containing media was removed and complete DMEM containing 10 µg/mL blasticidin (Thermo) was added. HepG2/C3A cells expressing dCas9-KRAB-MeCP2 were selected using 2 µg/mL puromycin (Sigma) and then maintained in culture with 1 µg/mL puromycin.

For western blot, cells were washed with HBSS for removing growth media and then incubated with lysis buffer (50 mM Tris-HCl ph 7.5, 0.1% SDS, 1% Triton X-100, 150 mM NaCl, and protease inhibitors) for 30 min on ice. Cell lysates were spun down for 5 min at 10,000 rpm at 4°C and the supernatant was transferred to a new tube. Protein concentrations were determined using a BCA protein assay kit (Thermo), according to manufacturer’s instructions.

### Western blot

Fifteen micrograms of cell lysate were subjected to SDS-PAGE using a 4 – 20% gradient gel, in the presence of β-mercaptoethanol (reducing conditions). For Western Blot, the proteins in the SDS-PAGE were transferred to a PVDF (polyvinylidene difluoride) membrane. The membrane was blocked with 5% non-fat dry milk in Tris-Buffered Saline (TBS)-Tween 0.1% for 1 h. The membrane was incubated with either anti-EXOSC4 (1:500, Santa Cruz) or anti-GAPDH (1:1,000, Santa Cruz) for 90 min. The membrane was washed with TBT-Tween 0.1% and immunoreactive proteins were detected using anti-mouse IgG conjugated with peroxidase (1:10,000, Jackson ImmunoResearch) for 1 h at room temperature. After washing five times for 6 min with TBS-Tween 0.1%, blots were developed using ECL (Amersham).

### Protein extraction and sample preparation

*Total proteome*: Cellular proteins were digested using S-Trap filters (Protifi), according to the manufacturer’s instructions. Briefly, cell pellets were resuspended with 5% SDS (sodium dodecyl sulfate), followed by incubation with 5 mM DTT (dithiothreitol, Sigma) for 1 h and with 20 mM iodoacetamide (Sigma) for 30 min, in order to reduce and alkylate proteins. Then, phosphoric acid was added to the samples at a final concentration of 1.2%. Proteins were diluted in six volumes of protein binding buffer (90% methanol and 10 mM ammonium bicarbonate, ph8), gentle mixed, and loaded to an S-Trap filter. Samples were spun at 500 g for 30 sec followed by two washes with binding buffer. One microgram of sequencing grade trypsin (Promega), diluted in 50 mM ammonium bicarbonate, was added to the S-Trap filter and proteins were digested overnight at 37°C. Peptides were eluted with 40 µl of 50 mM ammonium bicarbonate followed by 40 µl of TFA (trifluoroacetic acid) and 40 µl of 60% acetonitrile and 0.1% TFA. The peptide solution was pooled, spun at 1,000 g for 1 min, and dried in the SpeedVac.

### Nuclear proteome

The nuclear proteome was fractionated into soluble and chromatin-bound fractions. Briefly, the cell pellet was resuspended in five volumes of cold buffer A (10 mM ammonium bicarbonate pH 8, 1.5 mM MgCl_2_, and 10 mM KCl), incubated on ice for 10 min and centrifuged for 5 min at 400 g. The supernatant was removed, the pellet was resuspended in two volumes of buffer A containing 0.15% NP-40 and protease inhibitors and then centrifuged for 15 min at 3,200 g. The supernatant was stored as the cytoplasmic fraction. The pellet was washed with ten volumes of PBS, centrifuged for 5 min at 3,200 g, and the supernatant was discarded. The pellet was resuspended with two volumes of buffer C (420mM NaCl, 20mM ammonium bicarbonate pH 8.0, 20% glycerol, 2 mM MgCl_2_, 0.2 mM EDTA, 0.1% NP-40, 10 mM sodium butyrate, 0.5 mM DTT, and complete protease inhibitors) and incubated for 1 h at 4°C, on a rotating wheel. The suspension was centrifuged for 45 min at 20,800 g at 4°C. The supernatant was stored at - 80°C as the soluble fraction and the pellet as the chromatin-bound fraction.

### Histone extraction and sample preparation

Histone proteins were extracted from HepG2/C3A cell pellets, as described by (*52*). Briefly, histones were acid extracted with cold 0.2 M sulfuric acid (5:1, sulfuric acid:pellet) and incubated for 2 h at 4°C with constant rotation. Histones were then precipitated with 33% trichloroacetic acid (TCA) overnight at 4°C. The supernatant was removed, tubes were rinsed with ice-cold acetone containing 0.1% HCl, centrifuged and rinsed again with100% ice-cold acetone. The supernatant was discarded, and the pellet was dried using the SpeedVac. Before derivatization, histones were resuspended in 50 mM ammonium bicarbonate (pH 8) containing 20% acetonitrile. In the fume hood, 2 µl of propionic anhydride and 10 µl of ammonium hydroxide (all Sigma Aldrich) were added to the samples. The mixture was incubated for 5 min and the procedure was repeated. Histones were then digested with 1 µg of sequencing grade trypsin (Promega) (1:20, enzyme:sample) diluted in 50 mM ammonium bicarbonate, pH 8, and incubated overnight at 37°C. The derivatization reaction was repeated to derivatize peptide N-termini. The samples were dried in the Speedvac and stored in - 80°C.

### Pull-downs

*Peptide pull-down*: Synthesized biotinylated peptides were purchased from GenScript and resuspendend in HPLC water. Three peptides were used in this study: (i) K9me3, (ii) K14acK18acK23ac - 3 acetyls, and (iii) K9me3K14acK18acK23ac - hybrid. Twenty-five micrograms of peptide were incubated with 50 µl of Pierce Streptavidin Plus Ultra-Link Resin (Thermo) diluted in peptide binding buffer (150 mM NaCl, 50 mM Tris pH 8, and 0.1% NP-40), for 6 h at 4°C, on a rotating wheel. After incubation, the mixture of beads and peptide was washed three times with protein binding buffer (150 mM NaCl, 50 mM Tris pH 8, 10 µM ZnCl_2_, 0.5 mM DTT, 10 mM sodium butyrate, and protease inhibitors). Then, 100 µg of soluble nuclear proteins or chromatin-bound proteins, diluted in protein binding buffer, were incubated with the beads + peptides overnight at 4°C, on a rotating wheel. The mixture was spun briefly and the supernatant was discarded. The pellet was then washed five times with protein binding buffer containing 350 mM NaCl. Samples were reduced with 5 mM DTT (Sigma) for 1 h at room temperature and alkylated with 20 mM iodoacetamide (Sigma) for 30 min in the dark. Samples were spun and the supernatant was discarded. For digestion, samples were incubated overnight at 37°C with 500 ng sequencing grade modified trypsin (Promega). The supernatant containing the peptides were collected and dried using the SpeedVac.

### EXOSC4 pull-down

Twenty-five micrograms of nuclear, cytoplasmic or histone proteins, diluted in protein binding buffer, were incubated with 25 µg of EXOSC4 beads (Santa Cruz), overnight at 4°C, on a rotating wheel. As a negative control, proteins were incubated with 25 µg of streptavidin beads (Thermo). The mixture was spun briefly and the supernatant was discarded. The pellet was washed five times with protein binding buffer containing 350 mM NaCl. For proteome analysis, nuclear and cytoplasmic proteins were reduced with 5 mM DTT (Sigma) for 1 h at room temperature and alkylated with 20 mM iodoacetamide (Sigma) for 30 min in the dark. Samples were spun and the supernatant was discarded. For histone analysis, proteins were derivatized with propionic anhydride (Sigma) as described above. All samples were then digested overnight at 37°C with using 500 ng sequencing grade modified trypsin (Promega).

The supernatant containing the peptides were collected and dried using the SpeedVac.

### Sample desalting

Samples were desalted prior to mass spectrometry (MS) analysis as described by (*52*). Briefly, samples were resuspended in 100 µl of 0.1% TFA and loaded into a 96-well filter plate (Orochem) packed with Oasis HLB C-18 resin (Waters), which was equilibrated using 100 µl of the same buffer. Samples were washed with 100 µl of 0.1% TFA, eluted with 70 µl of a buffer containing 60% acetonitrile and 0.1% TFA, and dried using SpeedVac.

### Histone post-translational modifications (PTMs) analysis

After desalting, samples were resuspended in 10 µl of 0.1% TFA and loaded onto a Dionex RSLC Ultimate 300, coupled online with an Orbitrap Fusion Lumos (all Thermo Scientific). Chromatographic separation was performed using a two-column system, consisting of a C-18 trap cartridge (300 µm ID, 5 mm length) and an analytical column (75 µm ID, 25 cm length) packed in-house with reversed-phase Repro-Sil Pur C18-AQ 3 µm resin. Histone peptides were separated using a 60 min gradient from 4–30% buffer B (buffer A: 0.1% formic acid, buffer B: 80% acetonitrile + 0.1% formic acid) at a flow rate of 300 nl/min. The mass spectrometer was set to acquire spectra in a data-independent acquisition (DIA) mode. The full MS scan was set to 300–1100 m/z in the orbitrap with a resolution of 120,000 (at 200 m/z) and an AGC target of 5 × 105. MS/MS was performed in the orbitrap with sequential isolation windows of 50 m/z with an AGC target of 2 × 105 and an HCD collision energy of 30.

Raw files were imported into EpiProfile 2.0 software (*53*). From the extracted ion chromatogram, the area under the curve was obtained and used to estimate the abundance of each peptide. To achieve the relative abundance of PTMs, the sum of all different modified forms of a histone peptide was considered as 100% and the area of the particular peptide was divided by the total area for that histone peptide in all of its modified forms. The relative ratio of two isobaric forms was estimated by averaging the ratio for each fragment ion with different mass between the two species. The resulting peptide lists generated by EpiProfile were exported to Microsoft Excel and further processed for a detailed analysis.

### Proteomics analysis

After desalting, samples were resuspended in 10 µl of 0.1% TFA and loaded onto a Dionex RSLC Ultimate 300, coupled online with an Orbitrap Fusion Lumos (all Thermo Scientific). Chromatographic separation was performed using a two-column system, consisting of a C-18 trap cartridge (300 µm ID, 5 mm length) and an analytical column (75 µm ID, 25 cm length) packed in-house with reversed-phase Repro-Sil Pur C18-AQ 3 µm resin. Samples were separated using a 180 min gradient from 4 to 30% buffer B (buffer A: 0.1% formic acid, buffer B: 80% acetonitrile + 0.1% formic acid) at a flow rate of 300 nl/min. The mass spectrometer was set to acquire spectra in a data-dependent acquisition (DDA) mode. The full MS scan was set to 300–1200 m/z in the orbitrap with a resolution of 120,000 (at 200 m/z) and an AGC target of 5 × 10^5^. MS/MS was performed in the ion trap using the top speed mode (2 s), and AGC target of 1 × 10^4^ and an HCD collision energy of 35.

The MS raw data were processed using Proteome Discoverer software (v2.5, Thermo Scientific), SEQUEST search engine, and the SwissProt human database (updated June 2021). It was included variable modification of N-terminal acetylation and fixed modification of carbamidomethyl cysteine. Trypsin was specified as the digestive enzyme with two missed cleavages allowed. Mass tolerance was set to 10 ppm for precursor ions and 0.2 Da for product ions. Peptide and protein false discovery rate was set to 1%. Prior statistics, proteins were log2 transformed, normalized by the average value of each sample and missing values were imputed using a normal distribution 2 standard deviations lower than the mean as described (*54*).

Statistical regulation was assessed using heteroscedastic *T*-test (if *p*-value < 0.05). Data distribution was assumed to be normal, but this was not formally tested.

### RNA extraction and sequencing

One million cells per condition were stored in a DNA/RNA shield (Zymo) at -80°C. Cells were thawed on ice and the RNA was extracted using the RNeasy micro kit (Qiagen), according to the manufacturer’s instructions. RNA concentration and purity were measured using a Nanodrop (Thermo) and samples with a 260/280 ratio greater than 2 were further analyzed. RNA integrity number (RIN) was measured using a Bioanalyzer (Agilent) and samples with a RIN greater than 8 were kept. Samples were sent to Novogene for library preparation and whole transcriptome sequencing. Libraries were sequenced on the NovaSeq platform to obtain an average of 45 million paired-end 150 bp reads per sample.

### RNA-seq data analysis

Raw reads were trimmed to remove low quality base calls and Illumina universal adapters using Trim Galore! (Version 0.6.5) with default parameters and then assessed using fastQC (version 0.11.4) and multiqc (version 1.10.1). Reads were then aligned to the human genome (GRCh38) using STAR with default parameters. Alignment quality control was performed using RSeQC and Qualimap. Quantification was performed using RSEM. Quantification quality control was performed using EDASeq (version 2.3) and NOISeq (version 2.4). Gene ontology analysis was performed using clusterProfiler (version 4.4.4).

### Raw data availability

The mass spectrometry data have been deposited to the ProteomeXchange Consortium via the PRIDE (*55*) partner repository with the dataset identifier PXD053360.

## Acknowledgments

The Sidoli lab gratefully acknowledges for funding the Hevolution Foundation (AFAR), the Einstein-Mount Sinai Diabetes center, and the NIH Office of the Director (S10OD030286).

## Declaration of interests

The authors declare no competing interests.

## Notes

### Competing Interest Statement

The authors have declared no competing interest.

