## Supplementary data for "EXOSC4 is recruited by histone H3 co-modified with K9me3 and acetylations to surveil non-coding transcription"

*H3K9me3+acetyls surveil spurious transcription*

Stephanie Stransky[1], Ronald Cutler[1,2], Jennifer Aguilan[3], Joseph D. DeAngelo[1], David Shechter[1], Simone Sidoli[1*]

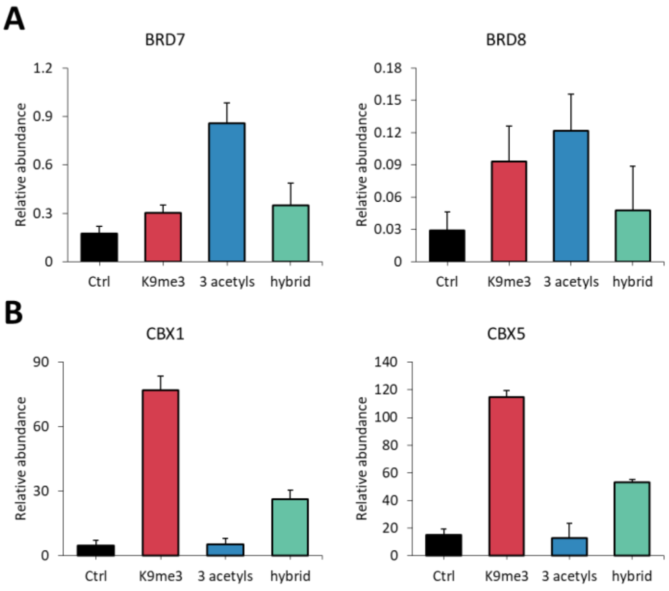


**Fig. S1. Peptide pull-down quality control.** Chromatin readers known for binding to specific histone post-translational modifications. **(A)** Bromodomains (BRD7 and 8) known to bind to hyperacetylated histones and **(B)** Chromodomains (CBX1 and 5) known to bind to histone methylation.

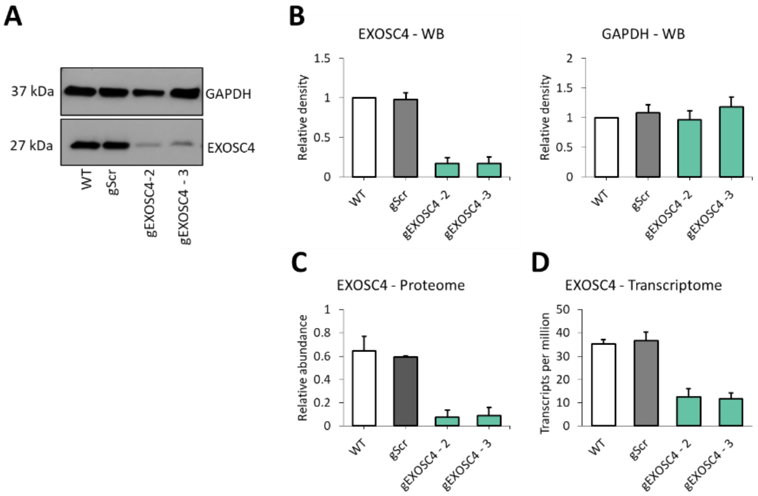


**Fig. S2. EXOSC4 knockdown. (A)** Western blot analysis of cell lysates after EXOSC4 KD. WT, wild type cells; gScr, scrambled guide RNA; gEXOSC4-2, EXOSC4 guide RNA 2; gEXOSC4-3, EXOSC4 guide RNA 3. **(B)** Quantification of the Western Blot bands. GAPDH was used as a loading control. **(C)** Proteome of EXOSC4 KD cells analyzed by mass spectrometry. **(D)** Transcriptome of EXOSC4 KD cells analyzed by RNA-seq. Data are represented as means ± SEM.

**Supplementary Table 1.** Histone analysis following 20mM sodium butyrate treatment.

**Supplementary Table 2.** Analysis of enriched proteins following peptide pull-down.

**Supplementary Table 3.** Analysis of enriched proteins following EXOSC4 pull-down.

**Supplementary Table 4.** Histone analysis following EXOSC4 pull-down.

**Supplementary Table 5.** List of gRNAs.
